# Bacterial metabolism rescues the inhibition of intestinal drug absorption by food and drug additives

**DOI:** 10.1101/821132

**Authors:** Ling Zou, Peter Spanogiannopoulos, Huan-Chieh Chien, Lindsey M. Pieper, Wenlong Cai, Natalia Khuri, Joshua Pottel, Bianca Vora, Zhanglin Ni, Eleftheria Tsakalozou, Wenjun Zhang, Brian K. Shoichet, Kathleen M. Giacomini, Peter J. Turnbaugh

## Abstract

Food and drugs contain diverse small molecule additives (excipients) with unclear impacts on human physiology. Here, we evaluate their potential impact on intestinal absorption, screening 136 unique compounds for inhibition of the key transporter OATP2B1. We identified and validated 24 potent OATP2B1 transport inhibitors, characterized by higher molecular weight and hydrophobicity compared to poor or non-inhibitors. OATP2B1 inhibitors were also enriched for dyes, including 8 azo (R−N=N−R′) dyes. Pharmacokinetic studies in mice confirmed that FD&C Red No. 40, a common azo dye excipient, inhibited drug absorption; however, the human gut microbiome inactivated azo dye excipients, producing metabolites that no longer inhibit OATP2B1 transport. These results support a beneficial role for the microbiome in limiting the unintended effects of food and drug additives in the intestine.

**One Sentence Summary:** Food and drug additives inhibit intestinal drug transporters, although some are inactivated by gut bacterial metabolism.

## Main text

One of the most notable aspects of life in the developed world is the routine exposure to chemicals through processed foods, pharmaceuticals, cosmetics, and the environment. Recent studies suggest that many of these small molecules have deleterious effects on human health (*1*), although the mechanisms through which they impact human pathophysiology remains poorly described. Even less is known about the reciprocal interactions between these compounds and the trillions of microorganisms that reside within the gastrointestinal tract (the gut microbiome) (*2*–*4*), despite emerging evidence that food and cosmetic additives can influence host-microbiome interactions (*1*, *5*).

In the current study, we focus on the bioactivity of pharmaceutical excipients — defined as substances other than the active pharmaceutical ingredients that are intentionally included in an approved drug delivery system or a finished drug product. On average, excipients make up 90% of a drug formulation and play crucial roles including assisting in stability, bioavailability, manufacturing, and patient acceptability (*6*). Excipients in oral drug products are present in intestinal fluid together with active ingredients; however, their impact on drug disposition is poorly understood. More broadly, many of the excipients used in pharmaceuticals are also widespread in processed food and cosmetics and therefore provides a routine source of chemical exposures for most individuals in the developed world.

We aimed to systematically screen the interactions between common pharmaceutical excipients and the intestinal transporter Organic Anion Transporting Polypeptide 2B1 (OATP2B1) by developing an *in vitro* assay to identify OATP2B1-inhibiting excipients. OATP2B1 (SLCO2B1) is localized to the apical membrane of the intestine and mediates the absorption of many oral prescription drugs, including the statins fluvastatin and rosuvastatin (*7*), the antihistamine fexofenadine (*8*, *9*), and the β-adrenoceptor blocker atenolol (*10*). Compounds that inhibit OATP2B1 transport may potentially reduce the absorption of many drugs. To identify excipients that interact with OATP2B1, we analyzed a comprehensive collection of molecular excipients identified using the Center of Excellence in Regulatory Science and Innovation (CERSI) Excipients Browser (*11*) and assembled a library of unique molecular excipients to screen for OATP2B1 transport inhibition. We excluded excipients that are no longer used, poorly soluble (solubility of <1 mM in H_2_O, DMSO, or ethanol), commercially unavailable, formulated for delivery by inhalation, or alternative salt forms of other excipients (e.g. sodium acetate was tested while calcium/magnesium acetate were not). This collection totaled 136 oral excipients spanning multiple functional classes including dyes, buffering agents, antimicrobial agents, and flavoring agents (**Fig. 1A**, **Fig. S1, Auxiliary Table 1**). This excipient collection was not exclusive to drugs, as 65% of our collection are also found in food products (**Auxiliary Table 1**).

**Fig. 1.**
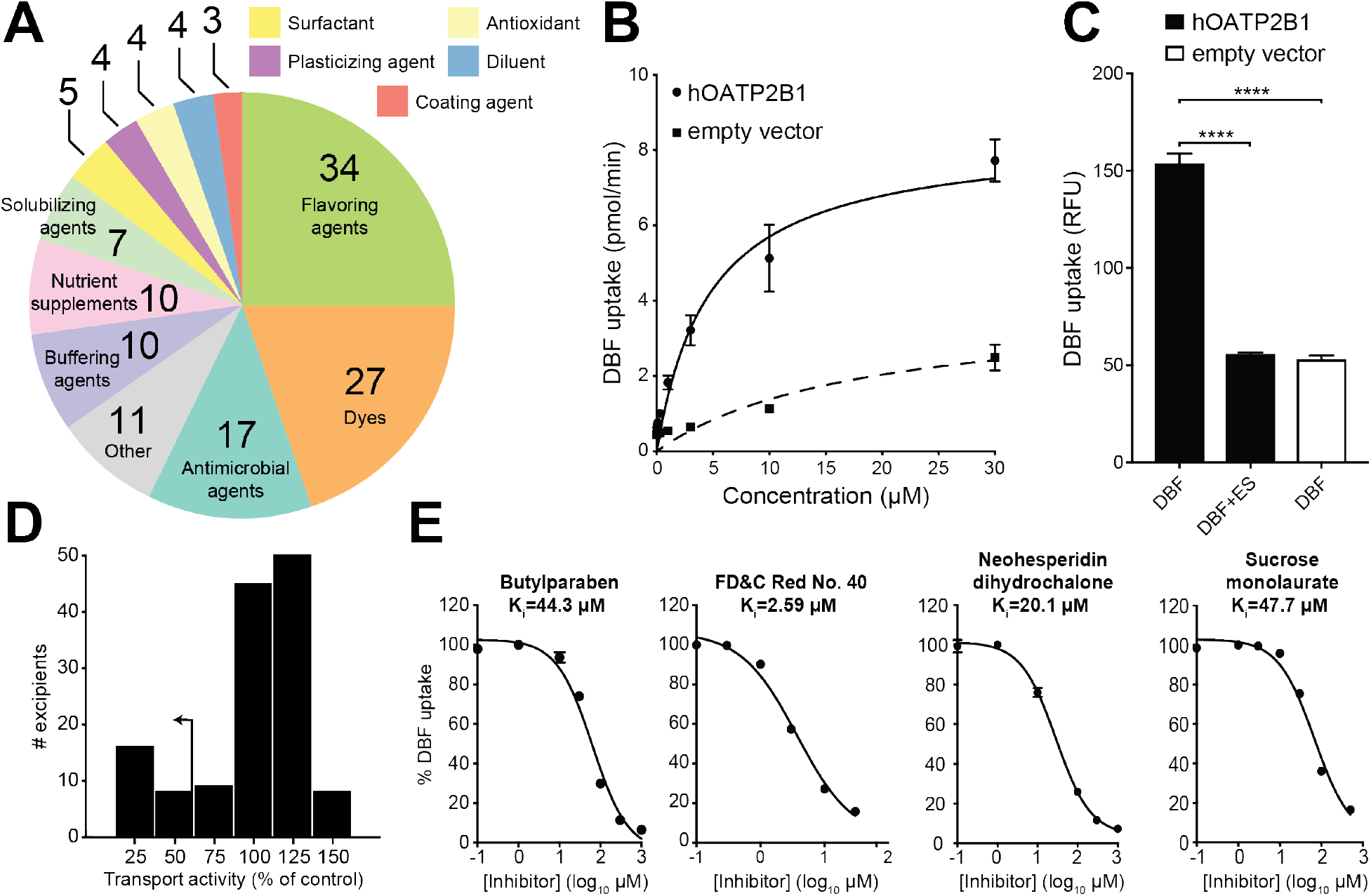
Multiple oral excipients inhibit OATP2B1-mediated transport. (A) Functional categories of the 136 oral molecular excipients included in our screen. (B) Kinetics of human OATP2B1-mediated 4’,5’-dibromofluorescein (DBF) uptake. Human OATP2B1-overexpressing (circles) and empty vector-transfected Human Embryonic Kidney (HEK) cells (squares) were incubated with DBF from 0.01 μM to 30 μM for 2 min. Data points represent the mean±SD of DBF uptake from ≥3 replicate determinations in three experiments. (C) Human OATP2B1-overexpressing HEK cells (black bars) were incubated in Hank’s Balanced Salt Solution (HBSS) uptake buffer containing 2 μM DBF for 3 min with or without 200 μM estrone sulfate (ES). Empty vector-transfected HEK cells (white bar) were assayed as above for background DBF uptake determination. Each column represents the mean±SD of DBF uptake from 8 replicate determinations. \*\*\*\**p*<0.0001, ANOVA with Dunnett’s correction. (D) Histogram showing the screening results of 136 excipients against OATP2B1. Our >50% inhibition cutoff is indicated by the arrow. (E) Dose-response curves of excipients against OATP2B1 transport. A representative excipient from each functional category is shown. Values represent the mean±SD of DBF uptake from three replicate determinations in a single experiment.

We developed an assay to identify potential inhibitors of OATP2B1-mediated uptake. The human *OATP2B1* cDNA was cloned into the pcDNA™ 5/FRT mammalian expression vector and transfected into human embryonic kidney (HEK) Flp-In cells to generate a stable OATP2B1-overexpressing cell line. The fluorescent molecule 4’,5’-dibromofluorescein (DBF) was used as a substrate of OATP2B1 uptake for screening. We determined a K_m_=4.7 μM for OATP2B1-mediated uptake of DBF (**Fig. 1B**), which is in agreement with the literature (*12*). The role of OATP2B1 in DBF uptake was confirmed by comparing our overexpressing cell line to our control cell line (HEK cells transfected with empty vector pcDNA™), revealing a significant (3-fold) increase in uptake under OATP2B1 overexpressing conditions (**Fig. 1C**). DBF uptake was significantly inhibited by the endogenous OATP2B1 substrate estrone sulfate (**Fig. 1C**). The statistical effect size (Z’-factor) was 0.79, indicating excellent assay quality (*13*). Together, these results demonstrate that our assay reliably and reproducibly measures the rate of OATP2B1-dependent DBF uptake.

Next, we used our validated assay to screen the full 136-member excipient library for inhibition of DBF uptake. Considering that the amount of excipients in an oral drug product can outweigh the active ingredient by 10- to 100-fold, we selected an excipient screening concentration of 200 μM (10-fold above the standard 20 μM used in drug screens (*14*); **Auxiliary Table 1**). We classified hits as excipients that inhibited OATP2B1 transport by >50% in order to focus on a manageable number of excipients for follow-up dose-response studies. Using this criterion, we identified 24 inhibitors in our primary screen (17.6% of the library; **Fig. 1D**). Next, we validated each of the 24 excipients by performing dose-response inhibition assays. This analysis validated 100% (24/24) of the inhibitors identified in our primary screen (**Fig. 1E**, **Table S1**), providing additional support for our >50% inhibition threshold and suggesting that additional excipient inhibitors remain to be discovered. Six excipients were potent inhibitors of OATP2B1-mediated uptake with an inhibition constant (K_i_) ≤1.0 μM (**Table S1**). We further validated two excipient inhibitors, FD&C Red No. 40 and FD&C Yellow No. 6, using the endogenous OATP2B1 substrate estrone sulfate (**Fig. S2**). An aggregation test on the 24 excipients demonstrated that the IC_50_ values of nine excipients were >10 times their aggregation concentration, suggesting specific inhibition of OATP2B1 (**Table S2**). These results indicate that the ability of oral drug excipients to inhibit drug uptake is far broader than previously appreciated (*15*).

The identified OATP2B1 transport inhibitors were chemically and functionally diverse, which included dyes, surfactants, antimicrobials, and flavoring agents (**Fig. 2A)**. Despite this diversity, we were able to identify characteristic signatures of OATP2B1 inhibitors relative to the overall library. Multiple physicochemical properties of excipient inhibitors were distinct relative to non-inhibitors, including increased molecular weight (*p*=3.80e-08, Student’s t-test; **Fig. 2B**) and lipophilicity (*p*=7.30e-11, Student’s t-test; **Figs. 2C and S3, Table S3**). We also found that OATP2B1 inhibitors were significantly enriched for dyes relative to the overall panel, representing 18/24 (75%) of the inhibitors (*p*<0.0001, Fisher’s exact test).

**Fig. 2.**
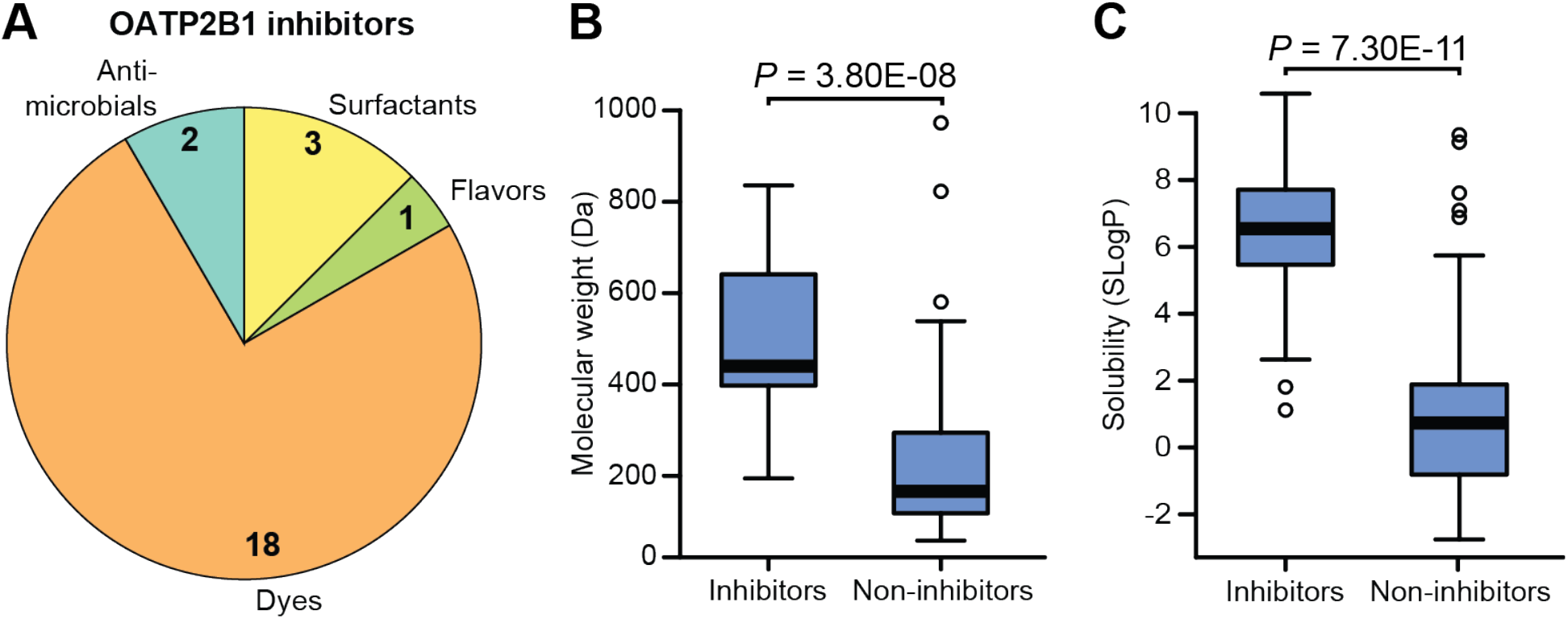
Excipient inhibitors of OATP2B1 transport have distinct physicochemical and functional properties compared to non-inhibitors. (A) Composition of the OATP2B1-inhibiting excipients identified from our screen. Excipient inhibitors were significantly enriched for dyes (*p*<0.0001, Fisher’s exact test). Excipient inhibitors display higher molecular weight (panel B) and solubility (SLogP) (panel C) compared to non-inhibitors. Box plots are shown for data in panels B, C (n=24 inhibitors versus 112 non-inhibitors) with the bold line representing the median. Points outside the bars are shown as open circles. *p*-values represent Student’s t-tests.

OATP2B1 plays an important role in fexofenadine absorption (*8*, *16*) and is a target of food-drug interactions (*9*, *17*). We selected the azo dye excipient FD&C Red No. 40 for *in vivo* studies because it is the dye with the highest approved amount in the United States by the Food and Drug Administration (FDA) and widely used in both food and drug products (*18*). The inhibitory effect of FD&C Red No. 40 on fexofenadine bioavailability was examined in P-glycoprotein (Pgp) deficient (*mdr1a/b*^−/−^) mice since fexofenadine is also a substrate of Pgp, which reduces fexofenadine absorption into systemic circulation (*19*). Concomitant administration of 25 mg/kg FD&C Red No. 40 (10 mM) significantly reduced fexofenadine area under the plasma concentration-time curve (AUC_0-360_) (n=9) by 48% compared to administration of the vehicle (n=8 mice/group, *p*=0.0026, ANOVA with Tukey’s correction; **Fig. 3, A and B**, **Table S4**). In contrast, administration of 2.5 mg/kg FD&C Red No. 40 (1 mM) (n=4) resulted in a fexofenadine AUC_0-360_ that was comparable to the control (*p*=0.8243).

**Fig. 3.**
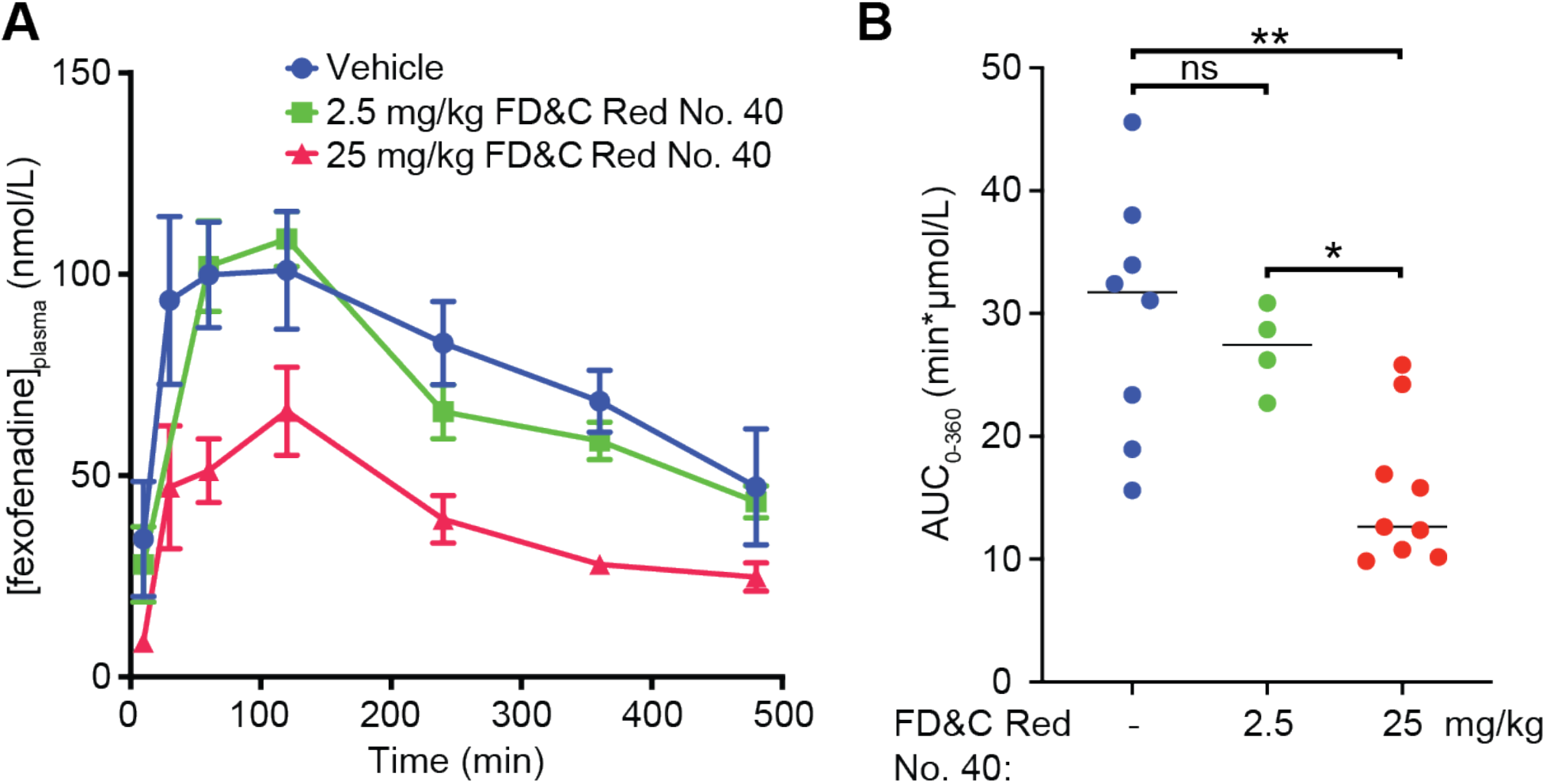
Oral administration of the OATP2B1 transport excipient inhibitor FD&C Red No. 40 reduces fexofenadine bioavailability in mice. (A) Plasma concentration-time profiles of fexofenadine after oral administration of 15 mg/kg fexofenadine with saline (blue circles, n=8), fexofenadine plus 2.5 mg/kg FD&C Red No. 40 (green squares, n=4), and fexofenadine plus 25 mg/kg FD&C Red No. 40 (red triangles, n=9) to P-glycoprotein (Pgp) deficient (*mdr1a/b*^−/−^) mice. Each point represents the mean±sem. (B) Plasma fexofenadine AUC_0-360_ calculated by Phoenix WinNonlin. Each point represents the plasma fexofenadine AUC_0-360_ from an individual mouse administered with 15 mg/kg fexofenadine plus saline (blue circles; n=8), fexofenadine plus 2.5 mg/kg FD&C Red No. 40 (green circles; n=4), and fexofenadine plus 25 mg/kg FD&C Red No. 40 (red circles; n=9). \**p*<0.05, \*\**p*<0.01; ANOVA with Tukey’s correction.

Of the 18 excipient dyes identified as inhibitors of OATP2B1 transport, eight belong to the azo dye family: synthetic dyes with one or more azo bonds (the functional group R−N=N−R′). The reductive cleavage of the azo bond is facilitated by azoreductases, which are encoded by phylogenetically diverse bacteria, including multiple bacterial taxa prevalent in the human gastrointestinal tract (*20*). Because azo dye excipients are orally administered, they have the opportunity of encountering and being cleaved by bacterial azoreductases, thus altering their chemical structure and potentially also their bioactivity.

Despite sharing an azo bond, azo dyes are structurally diverse and display variable susceptibility to reduction by bacteria (*21*, *22*). Furthermore, the ability of gut bacteria to metabolize the specific azo dyes identified in our screen was poorly understood (*21*, *23*, *24*). To test the ability of complex human gut microbiotas to metabolize the eight identified azo dye OATP2B1 inhibitors, we performed an *ex vivo* screen wherein each of the azo dye excipients were incubated with human stool samples from three unrelated healthy individuals. One dye was removed from this analysis for technical reasons. Following azo bond reduction, these dyes lose their chromogenic properties and therefore become colorless. All of the tested dyes were cleared by human gut bacteria (**Fig. 4A**). The extent of dye elimination varied between dyes, ranging from 90.5±2.9 to 48.2±7.3%, consistent with prior data suggesting that these enzymes display some degree of substrate specificity (*21*, *22*).

**Fig. 4.**
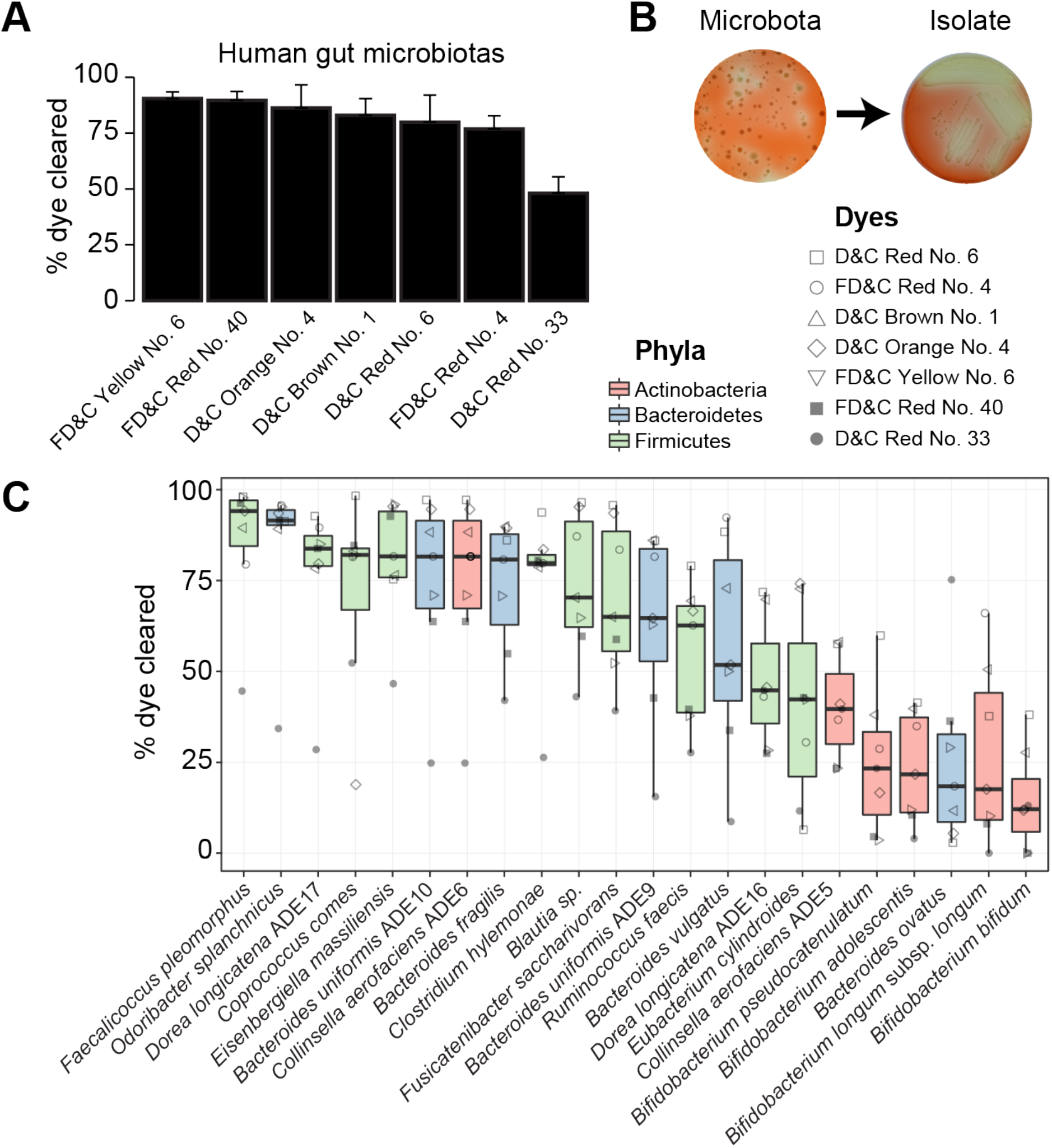
Human gut bacteria metabolize azo dye excipients. (A) Excipient azo dyes are metabolized following the *ex vivo* incubation with human fecal samples. Dilutions of human fecal suspensions from 3 healthy unrelated individuals were incubated anaerobically for 24 hours in BHI^+^ media supplemented with various excipient azo dyes and conditioned media was sampled and analyzed for residual dye concentration spectrophotometrically. Data is normalized to uninoculated controls. Values represent mean±sem. (B) Isolation of human gut bacteria capable of metabolizing FD&C Red No. 40. Dilutions of human fecal suspensions were used to inoculate BHI^+^ agar media supplemented with various excipient azo dyes and incubated anaerobically, with FD&C Red No. 40 shown here. Azo dye metabolizing colonies were identified by a zone of decolorization. Isolates capable of azo dye metabolism were picked and restreaked on BHI^+^ agar media supplemented with the same azo dye to confirm the phenotype. (C) Gut bacterial isolates show variable abilities to metabolize azo dye excipients. Human fecal bacterial isolates previously identified as azo dye metabolizers were incubated in triplicate in BHI^+^ media supplemented with various excipient azo dyes and incubated anaerobically for 24 hours. Residual dye was quantified spectrophotometrically from conditioned media samples and normalizing to uninoculated controls. Each point represents the mean dye clearance and the shape corresponds to the excipient azo dye. Colored boxes correspond to bacterial phyla.

Next, we developed a culture-based assay to identify human gut bacterial isolates capable of excipient azo dye metabolism. Dilutions of human fecal suspensions were plated on agar plates supplemented with azo dyes and incubated anaerobically. We identified metabolizers by inspecting agar plates for colonies that produced a zone of dye clearance, indicative of azo bond cleavage (**Figs. 4B and S4A**). Representative positive strains were selected and restreaked on azo dye containing media to confirm their phenotype (**Figs. 4B and S4, B and C**). First, we used a PCR-based fingerprinting method^33^ to dereplicate isolates from the same subject (*25*). Multiple instances were observed where the same bacterial fingerprint from a single human subject was observed across multiple dyes (**Auxiliary Table 2**).

To more definitively test for the ability of each isolate to metabolize multiple dyes we selected 22 unique azo dye excipient metabolizing bacterial isolates for 16S rRNA gene sequencing. We identified bacteria from the 3 major phyla found in the gut: Bacteroidetes (n=6), Firmicutes (n=10), and Actinobacteria (n=6) (**Auxiliary Table 2**). All of the bacterial strains tested were capable of clearing multiple azo dyes (**Fig. 4C**). The efficiency of dye elimination was primarily determined by strain not by dye (55% versus 19% of total variation; both factors *p*<0.0001, two-way ANOVA). This analysis also revealed a phylogenetic signature of azo dye excipient clearance, with bacteria belonging to the Firmicutes and Bacteroides phyla significantly more active than Actinobacteria (both comparisons *p*<0.0001, ANOVA with Tukey’s correction; **Figs. 4C and S5**). Together, these results indicate that human gut bacteria can metabolize multiple excipient dyes and that the extent of metabolism is influenced by bacterial taxonomy.

We hypothesized that bacterial metabolism of azo dye excipients would decrease their ability to inhibit OATP2B1 transport based on our previous observation that lower molecular weight is a characteristic feature distinguishing OATP2B1 inhibitors versus non-inhibitors (**Fig. 2B**). To test this hypothesis, we assayed conditioned media of human bacterial isolates grown in the presence of the azo dye FD&C Red No. 40 for OATP2B1 uptake inhibition. This representative azo dye excipient was selected due to our previous *in vivo* results (**Fig. 3**) and because it is the dye with the highest amount certified in the United States by the FDA and widely used in both food and drug products (*18*). Samples of conditioned media that showed complete clearance of FD&C Red No. 40 failed to inhibit OATP2B1-mediated [^3^H]-estrone sulfate uptake, whereas samples with high levels of FD&C Red No. 40 remaining inhibited uptake (**Fig. 5A**). LC-MS/MS analysis confirmed the depletion of FD&C Red No. 40 from conditioned media samples (**Fig. S6**). As expected, the level of clearance of FD&C Red No. 40 was significantly correlated with OATP2B1-mediated [^3^H]-estrone sulfate uptake (**Fig. 5B**).

**Fig. 5.**
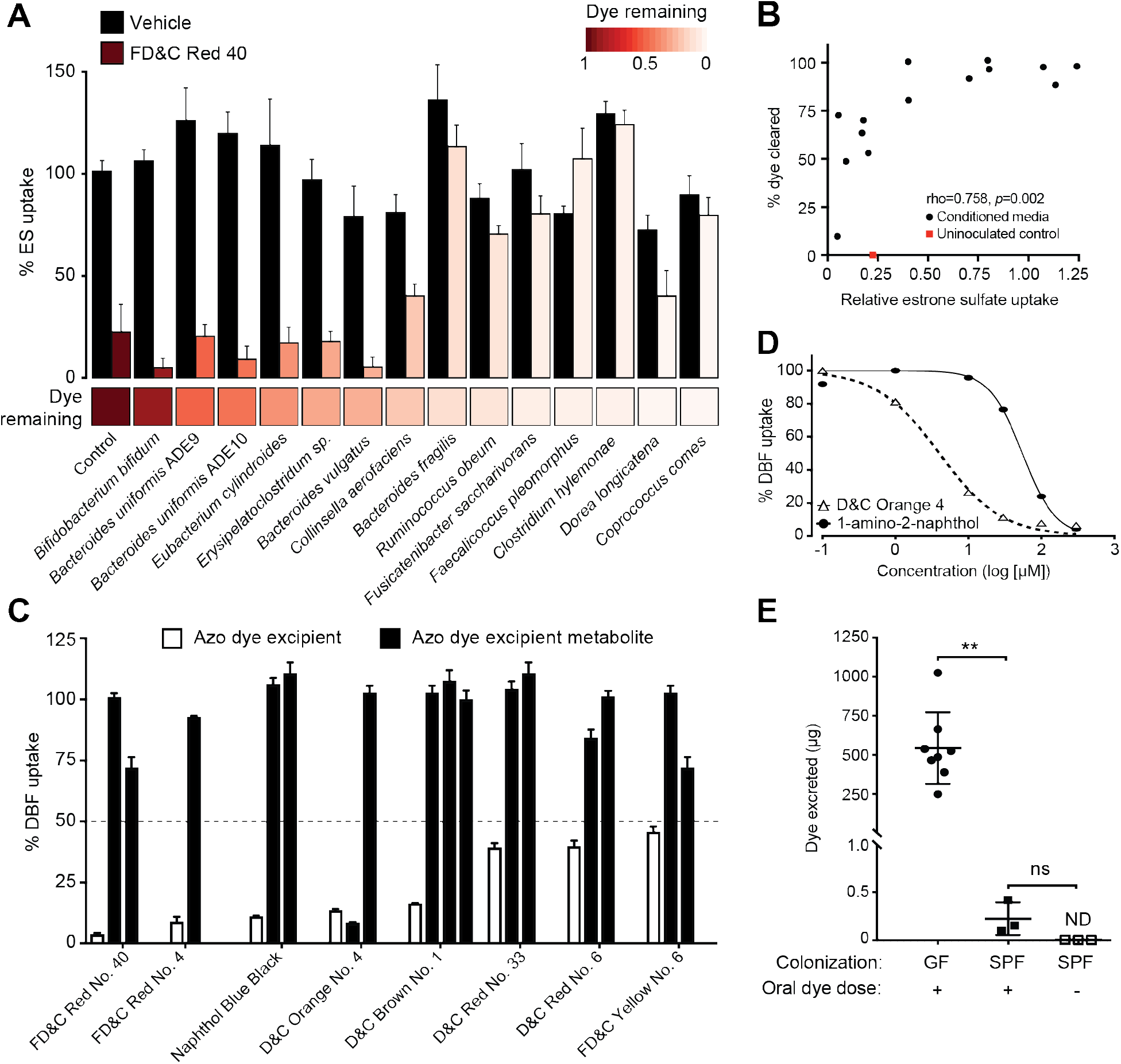
Microbial metabolism of azo dye excipients rescues OATP2B1 uptake inhibition. (A) Conditioned media samples of gut bacteria grown in the presence of 200 μM FD&C Red No. 40 or vehicle (DMSO) were assayed for the ability to inhibit OATP2B1 transport. Human OATP2B1-overexpressing cells were incubated with Hank’s Balanced Salt Solution (HBSS) uptake buffer containing 10 nM [^3^H]-estrone sulfate and bacterial conditioned media supplemented with FD&C Red No. 40 or vehicle (DMSO). Values represent the normalized mean±SD of [^3^H]-estrone sulfate uptake from three replicate determinations. The heat map below the bar graph shows relative FD&C Red No. 40 dye remaining in conditioned media samples, which was determined spectrophotometrically (B) FD&C Red No. 40 levels are associated with OATP2B1-mediated [^3^H]-estrone sulfate uptake (Spearman’s Correlation). (C) Inhibition of OATP2B1-mediated DBF uptake by azo dyes and their corresponding reduced metabolites. Human OATP2B1-overexpressing HEK cells were incubated in HBSS uptake buffer containing 2 μM DBF for 3 min with the designated concentrations of each azo dye and their metabolites. Concentrations assayed are specified in **Auxiliary Table 1** and **Table S5**. Values represent the mean±SD of normalized DBF uptake from n≥3 replicate determinations. (D) Dose-response curves of D&C Orange No. 4 and its metabolite, 1-amino-2-naphthol against OATP2B1-mediated DBF uptake. Values represent the mean±SD of DBF uptake from three replicate determinations. (E) Recovery of FD&C Red No. 40 dye in stool from germ-free and conventional mice. Mice were orally dosed with 5 mg of FD&C Red No. 40 and feces were collected over 24 hours and analyzed for residual dye levels. GF, germfree, SPF, conventional specific-pathogen-free. Values represent the mean±SD. \*\**p*<0.01, ANOVA with Tukey’s correction.

To more definitively test if this decrease in OATP2B1 inhibition was due to the biotransformation of azo dye excipients to their downstream microbial metabolites, we tested 12 unique metabolites from each of the 8 azo dye inhibitors. In contrast to the azo dye substrates, all of the corresponding reduced metabolites had a significant decrease in OATP2B1 transport inhibition (**Fig. 5C and Table S5)**. For example, the K_i_ value for D&C Orange No. 4 was 2.11 μM and the K_i_ values were 62.5 μM and >200 μM for 1-amino-2-naphthol and sulfanilic acid, respectively (**Fig. 5D**). This data highlighted that gut microbial metabolism rescues the inhibition of OATP2B1 transport by azo dye excipients.

Next, we administered FD&C Red No. 40 to germ-free and conventionally-raised mice to determine if gut microbial colonization alters the clearance of this excipient inhibitor following oral administration. Each animal was orally dosed with 5 mg of FD&C Red No. 40 and total feces were collected over 24 hours for residual azo dye analysis. Conventionally-raised (specific pathogen free, SPF) mice excreted significantly less FD&C Red No. 40 dye relative to germ-free controls (**Fig. 5E**). No dye was detected in untreated control SPF mice. These results suggest that the murine gut microbiome can efficiently metabolize azo dye excipients *in vivo* at the dose used; however, whether or not these excipients can impact OATP2B1 prior to their metabolism remains to be determined.

Finally, to assess the potential clinical relevance of the OATP2B1-inhibiting azo dye excipients, we estimated the maximal intestinal concentrations of azo dyes (**Table S6**). Based on the FDA perspective about the role of transporters in drug-drug interactions (*26*), it is possible that excipients with C_max_/K_i_≥10 could inhibit OATP2B1-mediated drug absorption at clinically relevant concentrations. However, it is important to point out that drug products frequently do not contain such large amount of these excipients, especially dyes. Larger amounts may be present in food products. It is worth noting that the amounts of artificial food colors certified for use in the United States by the FDA was 62 mg/capita/day in 2010, of which FD&C Red No. 40 accounts for 40%, FD&C Yellow No. 6 for 24%, FD&C Blue No.1 for 4%, and FD&C Red No. 3 for 4% (*18*). Based on 62 mg/day, the C_max_/K_i_ for FD&C Red No. 40 is 84.9 (**Table S6**). These data suggest that there may be food-drug interactions based on artificial food coloring agents inhibiting the OATP2B1-mediated absorption of prescription drugs.

Our results confirm and extend several recent *in vitro* studies suggesting that excipients can inhibit intestinal drug absorption (*27*–*29*). We discovered 24 excipients that are potent inhibitors of OATP2B1, 6 of which are predicted to achieve clinically-relevant intestinal concentrations (**Table S7**). Though clinical studies are needed to determine the *in vivo* effects of these excipients, these findings have potential clinical implications due to the key role of OATP2B1 in drug absorption (*9*, *30*, *31*). More broadly, pharmaceutical excipients constitute, on average, 90% of a drug formulation and yet they are often assumed to be inactive without being explicitly tested. These results suggest that pharmaceutical excipients harbor potentially unanticipated bioactivities, the downstream and clinical consequences of which remain poorly understood.

On the other hand, our data provide multiple potential strategies for mitigating the bioactivity of food and drug additives. These include characteristic physicochemical signatures of OATP2B1 inhibitors that could be used to shift the current formulation strategies from empirical to mechanistic-understanding-based formulation (*32*), which is of particular importance for the generic drug industry to develop bioequivalent drug product formulations. Furthermore, while our bacterial isolate data provide some support for the possibility that inter-individual differences in the gut microbiome could alter azo dye metabolism, the ubiquity and redundancy of this enzymatic activity between communities and taxonomic groups suggest that these specific excipients may have minimal intestinal activity in most human subjects.

In conclusion, these findings emphasize the importance of considering the chemical milieu in which drugs are taken and provide a mechanism through which the human gut microbiome ameliorates the impact of chemical exposures. This study also has clear translational implications, providing the first step towards the rational selection of excipients to minimize their off-target impacts on human and microbial cells.

## Supporting information

Auxiliary Table 1

Auxiliary Table 2

## Acknowledgements

Special thanks to Jordan Bisanz for comments on the manuscript. We are indebted to Jessie Turnbaugh in the UCSF Gnotobiotic Core for technical assistance. We thank Hayarpi Torosyan and Parnian Lak for their help and expertise in the aggregation assays.

## Funding

FDA (U01FD004979/U01FD005978) Office of Generic Drugs, which supports the UCSF-Stanford Center of Excellence in Regulatory Sciences and Innovation (UCSF-Stanford CERSI). National Institutes of Health (R01HL122593; R21CA227232) and the Searle Scholars Program (SSP-2016-1352). P.J.T. holds an Investigators in the Pathogenesis of Infectious Disease Award from the Burroughs Wellcome Fund, is a Chan Zuckerberg Biohub investigator, and is a Nadia’s Gift Foundation Innovator supported, in part, by the Damon Runyon Cancer Research Foundation (DRR-42-16). Fellowship support was provided by the Canadian Institutes of Health Research (P.S.) and the National Science Foundation (L.M.P.).

## Competing interests

P.J.T. is on the scientific advisory boards for Kaleido, Seres, SNIPRbiome, uBiome, and WholeBiome; there is no direct overlap between the current study and these consulting duties. All other authors have no relevant declarations.

## Data and materials availability

Data are available in the main text and supplementary materials. Figure source data and additional study data are available by request.

## Disclaimer

The contents of this publication are solely the responsibility of the authors and do not necessarily represent the official views of the Department of Health and Human Services or the US Food and Drug Administration (FDA).

## Supplementary Materials

Materials and Methods

Figs. S1-S6

Tables S1-S7

References (*34-44*)

Auxiliary Tables 1-2

## References and Notes

1. H. Yang, W. Wang, K. A. Romano, M. Gu, K. Z. Sanidad, D. Kim, J. Yang, B. Schmidt, D. Panigrahy, R. Pei, D. A. Martin, E. I. Ozay, Y. Wang, M. Song, B. W. Bolling, H. Xiao, L. M. Minter, G.-Y. Yang, Z. Liu, F. E. Rey, G. Zhang, A common antimicrobial additive increases colonic inflammation and colitis-associated colon tumorigenesis in mice. Sci. Transl. Med. 10(2018), doi:10.1126/scitranslmed.aan4116.

2. P. Spanogiannopoulos, E. N. Bess, R. N. Carmody, P. J. Turnbaugh, The microbial pharmacists within us: a metagenomic view of xenobiotic metabolism. Nat. Rev. Microbiol. 14, 273–287 (2016).

3. N. Koppel, V. Maini Rekdal, E. P. Balskus, Chemical transformation of xenobiotics by the human gut microbiota. Science. 356(2017), doi:10.1126/science.aag2770.

4. J. E. Bisanz, P. Spanogiannopoulos, L. M. Pieper, A. E. Bustion, P. J. Turnbaugh, How to Determine the Role of the Microbiome in Drug Disposition. Drug Metab. Dispos. 46, 1588–1595 (2018).

5. B. Chassaing, O. Koren, J. K. Goodrich, A. C. Poole, S. Srinivasan, R. E. Ley, A. T. Gewirtz, Dietary emulsifiers impact the mouse gut microbiota promoting colitis and metabolic syndrome. Nature. 519, 92–96 (2015).

6. Pharmaceutical excipients–where do we begin? NPS MedicineWise, (available at https://www.nps.org.au/australian-prescriber/articles/pharmaceutical-excipients-where-do-we-begin).

7. M. V. Varma, C. J. Rotter, J. Chupka, K. M. Whalen, D. B. Duignan, B. Feng, J. Litchfield, T. C. Goosen, A. F. El-Kattan, pH-sensitive interaction of HMG-CoA reductase inhibitors (statins) with organic anion transporting polypeptide 2B1. Mol. Pharm. 8, 1303–1313 (2011).

8. X. Ming, B. M. Knight, D. R. Thakker, Vectorial transport of fexofenadine across Caco-2 cells: involvement of apical uptake and basolateral efflux transporters. Mol. Pharm. 8, 1677–1686 (2011).

9. Y. Akamine, M. Miura, H. Komori, S. Saito, H. Kusuhara, I. Tamai, I. Ieiri, T. Uno, N. Yasui-Furukori, Effects of one-time apple juice ingestion on the pharmacokinetics of fexofenadine enantiomers. Eur. J. Clin. Pharmacol. 70, 1087–1095 (2014).

10. H. Jeon, I.-J. Jang, S. Lee, K. Ohashi, T. Kotegawa, I. Ieiri, J.-Y. Cho, S. H. Yoon, S.-G. Shin, K.-S. Yu, K. S. Lim, Apple juice greatly reduces systemic exposure to atenolol. Br. J. Clin. Pharmacol. 75, 172–179 (2013).

11. J. J. Irwin, J. Pottel, L. Zou, H. Wen, S. Zuk, X. Zhang, T. Sterling, B. K. Shoichet, R. Lionberger, K. M. Giacomini, A Molecular Basis for Innovation in Drug Excipients. Clin. Pharmacol. Ther. 101, 320–323 (2017).

12. S. Izumi, Y. Nozaki, T. Komori, O. Takenaka, K. Maeda, H. Kusuhara, Y. Sugiyama, Investigation of Fluorescein Derivatives as Substrates of Organic Anion Transporting Polypeptide (OATP) 1B1 To Develop Sensitive Fluorescence-Based OATP1B1 Inhibition Assays. Mol. Pharm. 13, 438–448 (2016).

13. J. H. Zhang, T. D. Chung, K. R. Oldenburg, A Simple Statistical Parameter for Use in Evaluation and Validation of High Throughput Screening Assays. J. Biomol. Screen. 4, 67–73 (1999).

14. N. Khuri, A. A. Zur, M. B. Wittwer, L. Lin, S. W. Yee, A. Sali, K. M. Giacomini, Computational Discovery and Experimental Validation of Inhibitors of the Human Intestinal Transporter OATP2B1. J. Chem. Inf. Model. 57, 1402–1413 (2017).

15. R. Panakanti, A. S. Narang, Impact of excipient interactions on drug bioavailability from solid dosage forms. Pharm. Res. 29, 2639–2659 (2012).

16. S. Medwid, M. M. J. Li, M. J. Knauer, K. Lin, S. E. Mansell, C. L. Schmerk, C. Zhu, K. E. Griffin, M. D. Yousif, G. K. Dresser, U. I. Schwarz, R. B. Kim, R. G. Tirona, Fexofenadine and Rosuvastatin Pharmacokinetics in Mice with Targeted Disruption of Organic Anion Transporting Polypeptide 2B1. Drug Metab. Dispos. 47, 832–842 (2019).

17. G. K. Dresser, D. G. Bailey, B. F. Leake, U. I. Schwarz, P. A. Dawson, D. J. Freeman, R. B. Kim, Fruit juices inhibit organic anion transporting polypeptide-mediated drug uptake to decrease the oral availability of fexofenadine. Clin. Pharmacol. Ther. 71, 11–20 (2002).

18. L. J. Stevens, T. Kuczek, J. R. Burgess, M. A. Stochelski, L. E. Arnold, L. Galland, Mechanisms of behavioral, atopic, and other reactions to artificial food colors in children. Nutr. Rev. 71, 268–281 (2013).

19. H. Tahara, H. Kusuhara, E. Fuse, Y. Sugiyama, P-glycoprotein plays a major role in the efflux of fexofenadine in the small intestine and blood-brain barrier, but only a limited role in its biliary excretion. Drug Metab. Dispos. 33, 963–968 (2005).

20. H. J. Haiser, P. J. Turnbaugh, Developing a metagenomic view of xenobiotic metabolism. Pharmacol. Res. 69, 21–31 (2013).

21. K. T. Chung, G. E. Fulk, M. Egan, Reduction of azo dyes by intestinal anaerobes. Appl. Environ. Microbiol. 35, 558–562 (1978).

22. K. T. Chung, S. E. Stevens Jr, C. E. Cerniglia, The reduction of azo dyes by the intestinal microflora. Crit. Rev. Microbiol. 18, 175–190 (1992).

23. R.-F. Wang, H. Chen, D. D. Paine, C. E. Cerniglia, Microarray method to monitor 40 intestinal bacterial species in the study of azo dye reduction. Biosens. Bioelectron. 20, 699–705 (2004).

24. F. Rafii, J. D. Hall, C. E. Cerniglia, Mutagenicity of azo dyes used in foods, drugs and cosmetics before and after reduction by Clostridium species from the human intestinal tract. Food Chem. Toxicol. 35, 897–901 (1997).

25. J. Versalovic, T. Koeuth, J. R. Lupski, Distribution of repetitive DNA sequences in eubacteria and application to fingerprinting of bacterial genomes. Nucleic Acids Res. 19, 6823–6831 (1991).

26. L. Zhang, Y. D. Zhang, J. M. Strong, K. S. Reynolds, S.-M. Huang, A regulatory viewpoint on transporter-based drug interactions. Xenobiotica. 38, 709–724 (2008).

27. N. Sjöstedt, F. Deng, O. Rauvala, T. Tepponen, H. Kidron, Interaction of Food Additives with Intestinal Efflux Transporters. Mol. Pharm. 14, 3824–3833 (2017).

28. R. Gurjar, C. Y. S. Chan, P. Curley, J. Sharp, J. Chiong, S. Rannard, M. Siccardi, A. Owen, Inhibitory Effects of Commonly Used Excipients on P-Glycoprotein in Vitro. Mol. Pharm. 15, 4835–4842 (2018).

29. A. Engel, S. Oswald, W. Siegmund, M. Keiser, Pharmaceutical excipients influence the function of human uptake transporting proteins. Mol. Pharm. 9, 2577–2581 (2012).

30. F. Qiang, B.-J. Lee, W. Lee, H.-K. Han, Pharmacokinetic drug interaction between fexofenadine and fluvastatin mediated by organic anion-transporting polypeptides in rats. Eur. J. Pharm. Sci. 37, 413–417 (2009).

31. T. Tapaninen, P. J. Neuvonen, M. Niemi, Orange and apple juice greatly reduce the plasma concentrations of the OATP2B1 substrate aliskiren. Br. J. Clin. Pharmacol. 71, 718–726 (2011).

32. R. A. Lionberger, FDA critical path initiatives: opportunities for generic drug development. AAPS J. 10, 103–109 (2008).

33. See Supplementary Materials and Methods.

