## Supplementary Materials for "Bacterial metabolism rescues the inhibition of intestinal drug absorption by food and drug additives"

**This PDF file includes:**

Materials and Methods
Figs. S1 to S6
Tables S1 to S7
Supplementary References

### Materials and Methods

Oral molecular excipient collection and reagents

As of March 2017, 292 oral molecular excipients listed on the UCSF-Stanford Center of Excellence in Regulatory Science Innovation (CERSI) Excipients Browser (http://excipients.ucsf.bkslab.org/excipients/molecular/?route=oral) were considered for screening (*34*). This collection was curated by excluding excipients that were duplicates, no longer used, having solubility issue, commercially unavailable, or formulated for delivery by inhalation. In total, 136 excipients were screened for OATP2B1-mediated transport inhibition (**Auxiliary Table 1**). 4′,5′-Dibromofluorescein (DBF) was purchased from Sigma-Aldrich (St. Louis, MO). Tritium labeled estrone sulfate was purchased from Perkin Elmer (Waltham, MA) with a specific activity of 54 Ci/mmol.

Establishment of stable OATP2B1-overexpressing cell line and cell culture

The human *OATP2B1* cDNA was cloned into pcDNA™ 5/FRT Mammalian Expression Vector (Thermo Fisher Scientific, Waltham, MA) and transfected into Flp-In™ 293 cells using Lipofectamine LTX (Life Technologies, Carlsbad, CA) according to the manufacturer's protocol. Stably transfected cells were maintained in Dulbecco's modified Eagle's medium (DMEM) supplemented with 10% fetal bovine serum, penicillin (100 U/ml), streptomycin (100 µg/ml), sodium pyruvate (110 µg/ml), and hygromycin B (100 µg/ml) at 37°C in a humidified incubator with 5% $CO_2$.

4′,5′-Dibromofluorescein uptake kinetics assay

The assay was based on a previously reported protocol (*35*) with minor modifications. Cells ($8 \times 10^4$/well) were seeded in black wall poly-D-lysine-coated 96-well plates 24 h prior to experiments. The DBF uptake kinetics study was conducted by incubating cells in Hank's Balanced Salt Solution (HBSS) containing 0.001 µM to 30 µM DBF at 37 °C for 3 min. Fluorescence in cells was measured using a fluorescence microplate reader with excitation and emission wavelengths at 485 nm and 560 nm, respectively. OATP2B1-mediated uptake of DBF was determined by subtracting background uptake from the empty vector-transfected cells at each substrate concentration. The Z' assay factor was calculated according to the equation: $Z' = 1 - 3 \times (\delta_{control+} + \delta_{control-})/(\mu_{control+} - \mu_{control-})$, where δ and µ are the standard deviation and the mean, respectively (*36*).

Excipient screening and $IC_{50}$ determination

A total of 136 excipients were screened at 200 µM with 1% (v/v) of vehicle (DMSO), except for several surfactants and those with limited solubility, which were tested at lower concentrations (**Auxiliary Table 2**). Downstream reduced metabolites of selected azo dye excipients were synthesized (Advinus Therapeutics, Pune, India) and tested at 200 µM, except for those with limited solubility, which were tested at lower concentrations (**Table S5**). The screen was performed in triplicate with 2 µM DBF. The estrone sulfate uptake assay was conducted by incubating cells in HBSS containing 0.02 µM [$^3$H]-estrone sulfate for three minutes at 37 °C. The cells were then washed twice with ice-cold HBSS followed by lysis using lysis buffer (0.1 N NaOH and 0.1% SDS). [$^3$H]-estrone sulfate uptake was determine by radioactivity in the lysate using a liquid scintillation counter. $IC_{50}$ values for compounds were determined by fitting the uptake results to the Hill equation by non-linear regression using GraphPad Prism. The inhibition constant ($K_i$) was estimated from the following equation (*37*):

$$Ki = \frac{IC50}{1 + \frac{S}{Km}} \quad (1)$$

The $K_m$ value of DBF for OATP2B1 is 4.7 μM and S is 2 μM.

Estimation of maximum intestinal concentrations of excipients

The Accepted Daily Intake (ADI) or maximum allowable amount of each excipient added for oral consumption was used to estimate the maximum intestinal concentration ($C_{max}$) as shown in the following equation. The FDA perspective about the role of transporters in drug-drug interactions indicates that drugs exhibiting $[I]/IC_{50}$ (or $K_i$) ≥10 should be evaluated for potential clinical drug-drug interactions (*38*). [I] is the theoretical maximal intestinal concentration of a drug after oral administration calculated as the highest clinical dose (mg) in a volume of 250 ml.

$$Assumed\ maximum\ intestinal\ concentration\ (Cmax) = \frac{Amount\ (g) \div Molar\ Mass\ (g/mol)}{250\ mL} \quad (2)$$

Aggregation test

The test for aggregation was performed according to a previously reported protocol (*39*). Dynamic light scattering experiments were carried out in a 384-well plate using a modified DynaPro Plate Reader II system (Wyatt Technology); the width of the 60 mW laser beam (at ~830 nm) was expanded to be appropriate for detecting large colloidal particles. Version 1.7 of the Dynamics software (Wyatt Technology) was used to process the data, as well as to automatically adjust laser power and detector angle (158 degrees). The compounds were each concentrated in DMSO, subsequently diluted with filtered 50 mM potassium phosphate, pH 7.0, leading to a final concentration of 1% DMSO, and then added to the plates for measurement.

Analysis of physicochemical properties of OATP2B1

Molecular structure files in structure-data format (SDF) were obtained from the CERSI Excipient Browser at http://excipients.ucsf.bkslab.org/ and the PubChem database (*40*). MayaChemTools software (*41*) was used to compute eight one-dimensional and two-dimensional descriptors, such as molecular weight (MW), number of heavy atoms (HA), number of rotatable bonds (RB), molecular volume (MV), number of hydrogen bond donors (HBD), number of hydrogen bond acceptors (HBA), Crippen's log of the octanol/water partition coefficient (including implicit hydrogens) (SLogP), and total polar surface area (TPSA). Boxplots were prepared using R software. Differences between distributions of molecular descriptors were assessed by the Student's t-test using R software. The R package FSelector (*42*) was used to select two non-redundant and informative physicochemical properties from the eight descriptors (e.g. molecular weight and SLogP).

Fexofenadine pharmacokinetics in Pgp deficient mice

Mouse experiments were approved by the University of California, San Francisco Institutional Animal Care and Use Committee. 8-10 week-old male P-glycoprotein (Pgp) deficient (*mdr1a/b*$^{-/-}$

) mice (Taconic Biosciences, Model#: 1487-M) were fasted overnight prior to drug and excipient oral administration. For the control group, saline was administered by oral gavage immediately followed by 15 mg/kg fexofenadine by oral administration. For test groups, FD&C Red No. 40 was administered to at a concentration of 2.5 mg/kg (1 mM) or 25 mg/kg (10 mM) by oral gavage immediately followed by 15 mg/kg fexofenadine by oral administration. Blood samples (~25 µL each) were obtained from mouse tail vein at 10, 30, 60, 120, 240, 360, and 480 min post drug administration in heparinized glass micro-hematocrit capillary tubes. Blood samples were centrifuged at 3500xg for 15 min, plasma was recovered and stored at -80°C until analysis. Pharmacokinetic parameters were computed via noncompartmental analysis using Phoenix WinNonlin v8.1 (Certara, Princeton, NJ). Area under the curve (AUC) was calculated using the linear-up, log-down method. Exact sample collection times were used for pharmacokinetic analysis.

Liquid Chromatography-Mass Spectrometry analysis of plasma fexofenadine

Four times the volume of acetonitrile was added to plasma samples, vigorously vortexed, and centrifuged at 12,000 rpm for 5 min at room temperature. The supernatant was recovered and used for analysis. LC-HRMS analysis was performed using an Agilent Technologies 6520 Accurate-Mass Q-TOF LC-MS instrument and an Agilent Eclipse Plus C18 column (4.6 x 100 mm). A linear gradient of 2-98% acetonitrile (v/v) over 30 min in $H_2O$ with 0.1% formic acid (v/v) at a flow rate of 0.5 mL/min was used. Data was acquired under positive mode and 20 ppm mass error tolerance was used for each trace.

Screening human stool samples and bacterial isolates for azo dye metabolism in liquid cultures

All experiments were performed under the guidance of the UCSF Institutional Review Board. Following informed consent, a single stool sample was collected from three healthy adult volunteers and frozen immediately at -80°C. On the day of the assay for bacterial dye metabolism, the sample was thawed on ice. All downstream procedures and incubations were performed in an anaerobic chamber (COY Laboratory Products Inc.) with the following atmosphere: 10% $H_2$, 5% $CO_2$, and 85% $N_2$. All reagents were equilibrated in the anaerobic chamber for at least 24 hours before using. Three independent segments of a single stool specimen were sampled and added to supplemented Brain-Heart Infusion ($BHI^+$) medium at a ratio of 15 ml/g wet weight. $BHI^+$ is composed of: L-cysteine hydrochloride (0.05% w/v), hemin (5 µg/mL), and vitamin K (1 µg/mL). Samples were vortexed vigorously for 5 min and then allowed to settle for 10 min. Fecal suspensions were then diluted 10-fold serially in $BHI^+$ medium and 70 µl the $10^{-3}$ dilution was used to inoculate Hungate tubes with 7 ml of $BHI^+$ medium supplemented with azo dye at a final concentration of 250 µM. Azo dye stocks were prepared in DMSO at a concentration of 25 mM and used at final concatenation of 1% (v/v). Naphthol blue black was excluded due to solubility issues. Cultures were incubated at 37°C. A conditioned media sample was taken at 24 hours, centrifuged to remove cells, and analyzed for residual dye concentration by measuring absorbance at 450 nm ($A_{450\ nm}$). A sterile control with no added dye ($BHI^+$ medium only) was used as the background $A_{450\ nm}$ and subtracted from each sample. Percent dye metabolized was calculated by dividing $A_{450\ nm}$ from conditioned media samples by the $A_{450\ nm}$ of the sterile control supplemented with dye (uninoculated $BHI^+$ medium plus azo dye) and multiplying by 100. Human fecal bacterial isolates were screened in a 96-deep-well plate using the same media and dye concentrations described above. Overnight cultures of bacterial isolates were grown in Hungate tubes and adjusted to an $OD_{600\ nm}$ of 0.1. A 15 µl aliquot of this bacterial suspension was used to inoculate 1.5 ml of dye containing media and cultures were incubated at 37°C for 24 hours.

Isolating human gut bacteria capable of azo dye metabolism

Human fecal suspensions were prepared as described above. $BHI^+$ agar medium supplemented with 250 µM dyes were inoculated with 1 ml of a $10^{-6}$ fecal suspension dilution prepared as described above. The human fecal samples ranged from 1-5 × $10^9$ CFUs/ml/g of fecal sample. Agar plates were incubated anaerobically at 37°C for up to a week. Azo dye reducing frequency was calculated by dividing the number of metabolizing colonies by the total number of colonies on each plate. Colonies that showed dye metabolism (decolorized halo) were picked and restreaked on media supplemented with dye to confirm the phenotype.

Dereplication of human bacterial isolates and 16S rRNA gene amplification and sequencing

Genomic DNA from each strain was prepared using the Qiagen DNeasy Blood & Tissue Kit (Hilden, Germany) following the manufacturer's protocol for bacteria. Dereplication of isolated bacterial strains was accomplished using Enterobacterial repetitive intergenic consensus PCR (ERIC-PCR) with the following primers: ERIC2, 5'-AAGTAAGTGACTGGGGTGAGCG-3', and ERIC1R, 5'-ATGTAAGCTCCTGGGGATTCAC-3'(*43*). Strains with identical ERIC-PCR product fingerprints were considered replicates. The 16S rRNA gene was amplified from representative isolates using Thermo Fisher Scientific AmpliTaq Gold DNA polymerase (South San Francisco, CA) with the following primers: 8F, 5'- AGAGTTTGATCCTGGCTCAG-3'; 1391R, 5'- GACGGGCGGTGWGTRCA-3' (*44*). Sanger sequencing of PCR products was performed by GENEWIZ. Species were determined by comparing 16S rRNA gene sequences to the NCBI nucleotide collection database.

Assaying conditioned azo dye containing media for inhibition of OATP2B1 uptake

Human gut bacterial isolates were grown as described above section *Screening human stool samples and bacterial isolates for azo dye metabolism in liquid cultured*. Following 48 hours of incubation, 96-well plates were centrifuged to pellet bacterial cells and the resulting conditioned media was recovered. Inhibition of OATP2B1-mediated [$^3$H]-estrone sulfate uptake in OATP2B1-transfected HEK cell was performed as described in the above section *Excipient screening and $IC_{50}$ determination* with the addition of conditioned media samples at 10% (v/v).

Liquid Chromatography-Mass Spectrometry analysis of FD&C Red No. 40 from bacterial conditioned media samples

Bacteria cultures grown in the presence of FD&C Red No. 40 were pelleted by centrifugation (4,000xg for 15 min), and the conditioned media was extracted 1:1 by volume with ethyl acetate. The ethyl acetate was removed by rotary evaporation, and the dry extract was dissolved in 100 µL of methanol. LC-HRMS analysis was performed using Agilent G6430 QQQ instrument and an Agilent Eclipse Plus C18 column (4.6 x 100 mm). A linear gradient of 2-98% acetonitrile (v/v) over 30 min in $H_2O$ with 0.1% formic acid (v/v) at a flow rate of 0.5 mL/min was used. Data was acquired under negative mode and 20 ppm mass error tolerance was used for each trace.

Quantification of FD&C Red No. 40 from murine fecal samples

Mouse experiments were approved by the University of California, San Francisco Institutional Animal Care and Use Committee and conducted with support from the UCSF Gnotobiotic Core Facility. Conventional animals were housed in a specific animal free (SPF) facility. Female C57BL/6 mice (Taconic Biosciences, Model#: B6-F) were used between 8-10 weeks of age. Germ-free female C57BL/6J mice between 8-10 weeks of age were bred in-house in sterile breeding isolators (Class Biologically Clean, USA) and transferred to experimental isolators prior ro FD&C Red No. 40 dosing. Mice were fed a standard autoclaved chow diet *ad libitum* (Lab Diet 5021) with free access to water at room temperature (24 ± 1°C) and a 12/12-h light/dark cycle (7:00 a.m.–7:00 p.m.). Mice were orally dosed (0.2 mL) with 5 mg of FD&C Red No. 40 dissolved in sterile water and immediately individually housed in a metabolic chamber for mouse (Harvard Apparatus, 72-7060) for 24 hours. A control group of mice conventional SPF mice were dosed with sterile water instead of FD&C Red No. 40. Fecal pellets were collected and stored at -80°C. Fecal pellets were lyophilized overnight to dryness and accurately weighed. 300 μL of methanol was added and FD&C Red No. 40 was extracted for 40 min by sonication at room temperature. The supernatant was analyzed by LC-MS as described above.

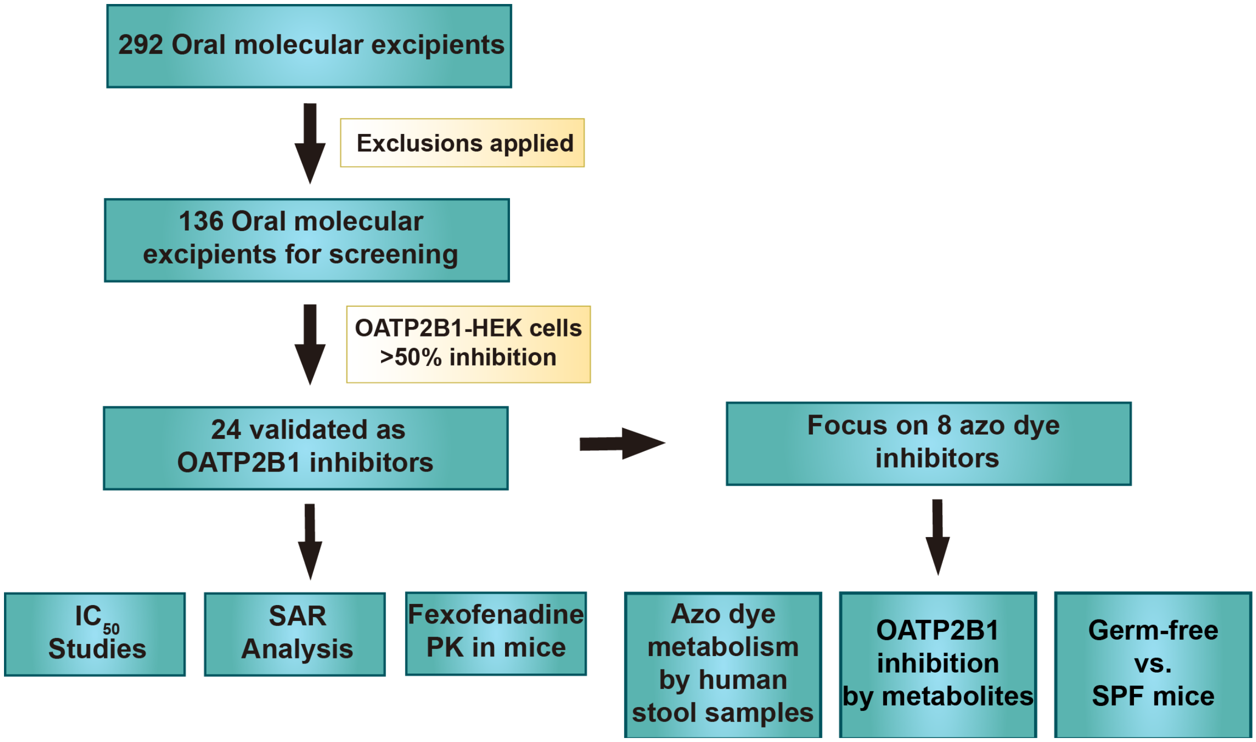


**Fig. S1. Overview of our study design.** In total 136 oral molecular excipients were selected and screened for inhibition of OATP2B1-mediated DBF uptake in OATP2B1-overexpressing HEK293 cells. $IC_{50}$ studies, quantitative structure-activity relationship (SAR) were conducted on 24 identified inhibitors and fexofenadine pharmacokinetics (PK) were evaluated to examine the inhibitory effect of the representative OATP2B1 inhibitor, FD&C Red No. 40, in Pgp-deficient (*mdr1a/b*$^{-/-}$) mice. In addition, the metabolism of OATP2B1-inhibiting azo dyes by gut bacteria was assessed through *ex vivo* incubations of healthy human stool samples, *in vitro* analysis of gut bacterial isolates, and experiments in germ-free and conventionally-raised (SPF, specific pathogen free) mice. The inhibitory effect of azo dyes and their bacterial metabolites on OATP2B1 was also determined.

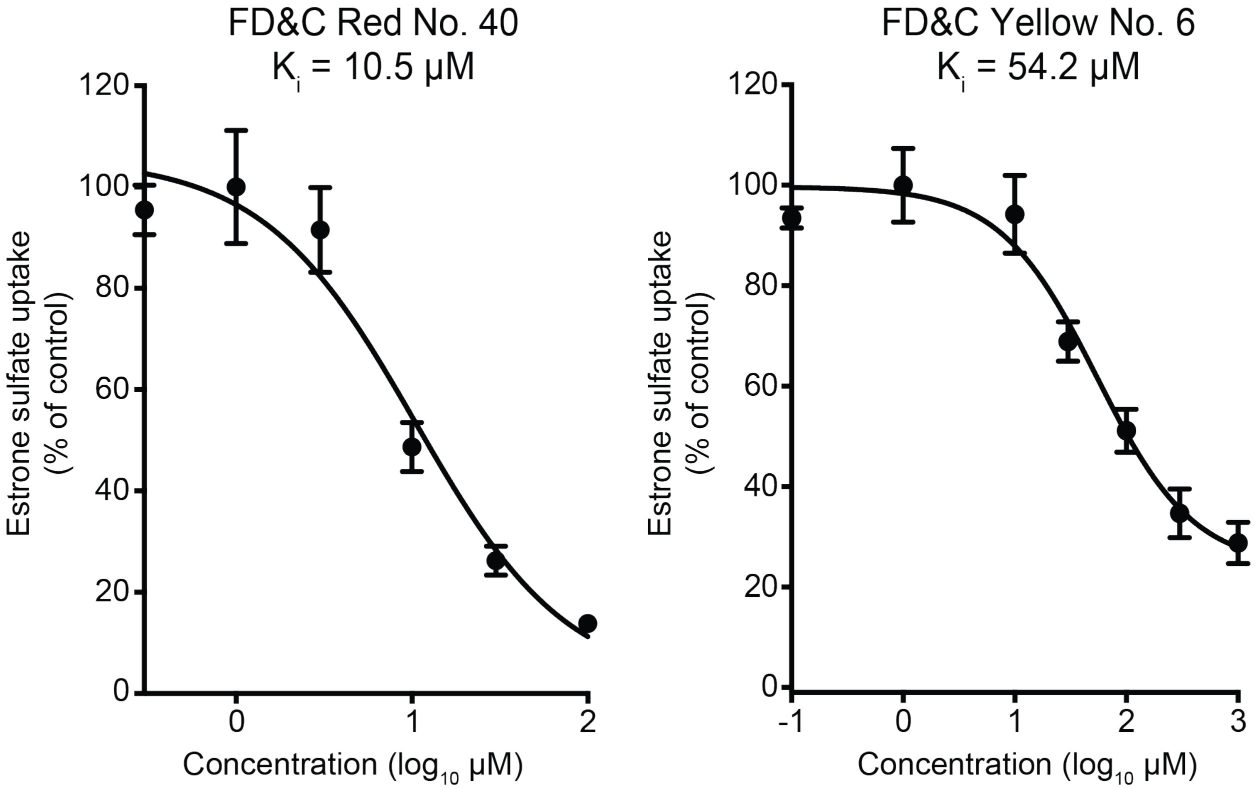


**Fig. S2. Dose-dependent inhibition of OATP2B1-mediated [$^3$H]-estrone sulfate (ES) uptake by FD&C Red No. 40 and FD&C Yellow No. 6.** Human OATP2B1-overexpressing cells were incubated with Hank's Balanced Salt Solution (HBSS) uptake buffer containing 10 nM [$^3$H]-estrone sulfate with designated concentrations of FD&C Red No. 40 or FD&C Yellow No. 6 for 3 min. Data points represent the mean±SD of [$^3$H]-estrone sulfate uptake from three replicate determinations.

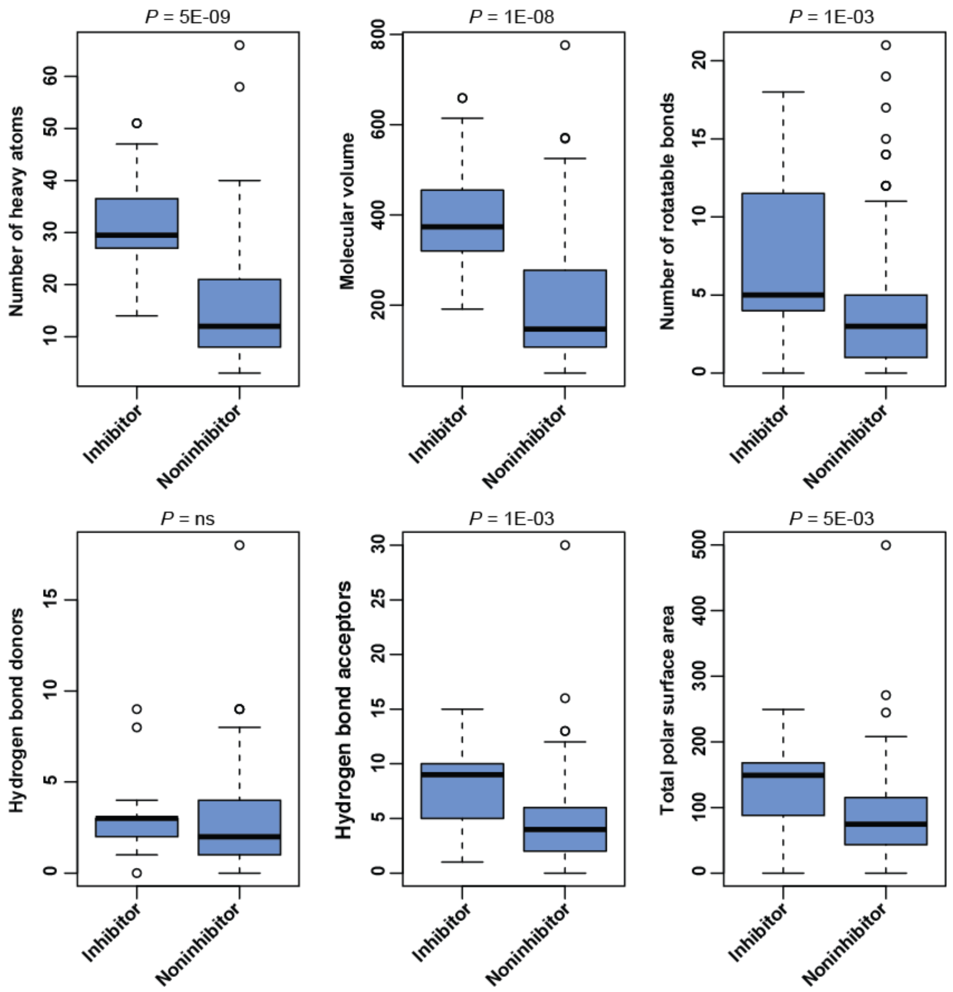


**Fig. S3. Comparison of physicochemical properties between excipient inhibitors and non-inhibitors of OATP2B1.** Box plots are shown (n=24 inhibitors versus 112 non-inhibitors) with the bold line representing the median and the error bars the upper and bottom quartiles. Points outside the bars are shown as open circles. *p*-values represent Student's t-tests.

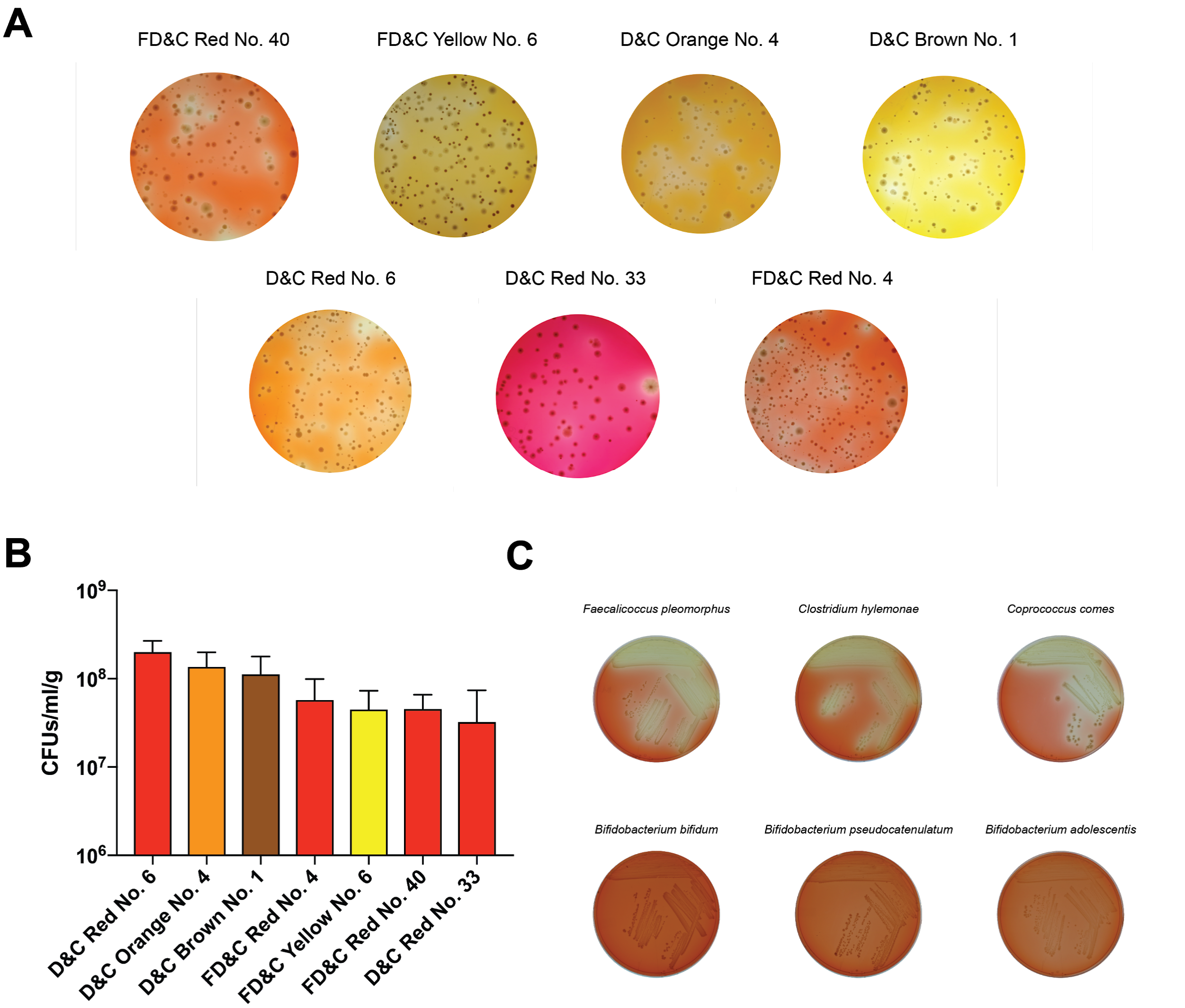


**Fig. S4. Isolation of azo dye excipient metabolizing human gut bacterial isolates.** (A) A dilution of human fecal suspensions were plated on $BHI^+$ media supplemented with excipient azo dyes and incubated anaerobically. Metabolizing colonies were identified by a characteristic decolorization halo surrounding the colony. (B) Recovery numbers of azo dye metabolizing bacterial isolates from 3 healthy human fecal samples. Bar plots show mean ±SD. (C) Azo dye metabolizing colonies were restreaked on $BHI^+$ media supplemented with azo dye to confirm their phenotype. Top agar plates show isolates capable of FD&C Red No. 40 metabolism. The bottom plates show isolates that cannot metabolize FD&C Red No. 40.

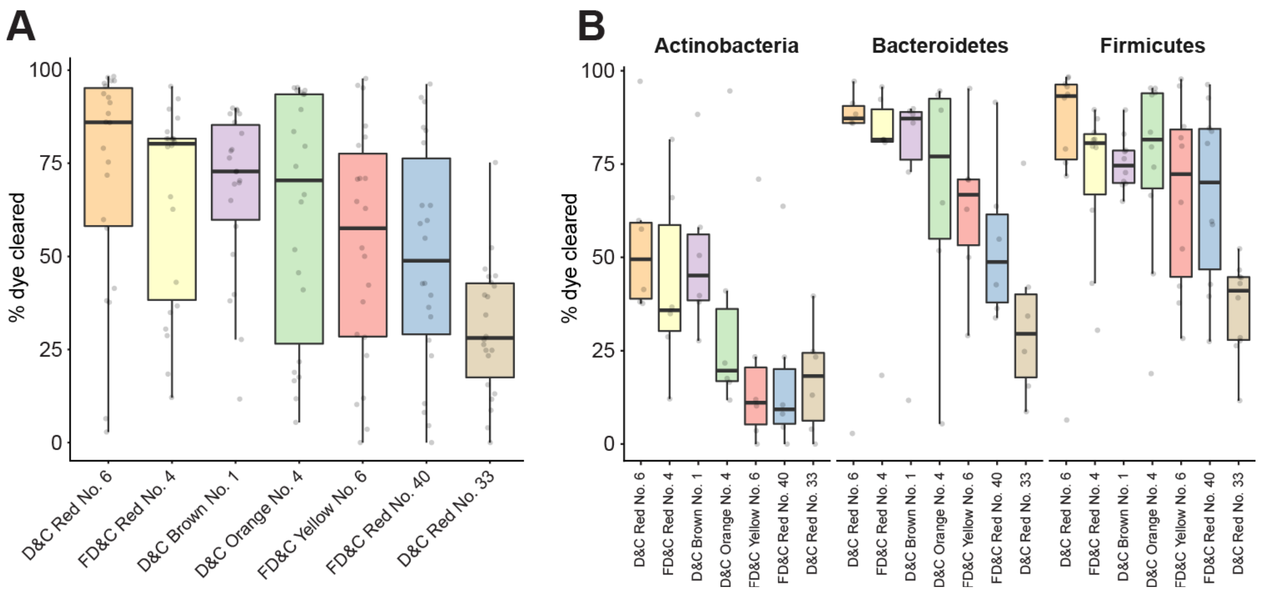


**Fig. S5. Metabolism of azo dye excipients by human gut bacterial isolates.** 22 unique human gut bacterial isolates recovered from 3 healthy human fecal samples were assayed for metabolism against 7 azo dye excipients. Bacteria isolates were inoculated in BHI$^{+}$ media supplemented with azo dye and incubated anaerobically for 24 hours. Conditioned media was recovered and assayed for residual azo dye spectrophotometrically. All assays were performed in triplicate and normalized to an uninoculated control sample. (A) Boxplots displaying percent azo dye metabolized by all strains for each azo dye. Each dot represents the mean metabolism by a single bacterial isolate. (B) Boxplots displaying percent azo dye metabolized grouped according to phylum for each azo dye. Each dot represents the mean metabolism by a single bacterial isolate.

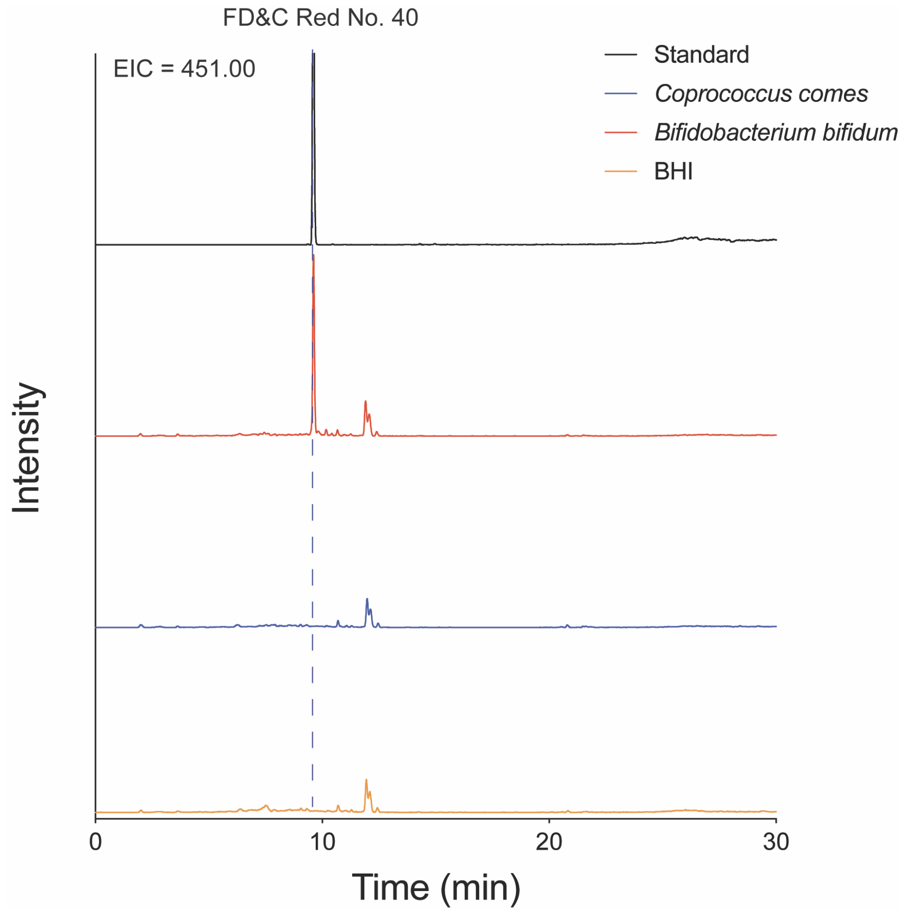


**Fig. S6. Measurement of residual FD&C Red No. 40 in conditioned media samples by LC/MS.** Extracted ion chromatograms (EICs) show the presence of FD&C Red No. 40 in the conditioned media sample of a non-metabolizing isolate (*Bifidobacterium bifidum*, red), but could not be detected in a sample from a metabolizing isolate (*Coprococcus comes*, blue). Sterile media control is shown in yellow. A 20 ppm error was used for each trace.

**Table S1. Validation and inhibitory potencies for 24 OATP2B1 inhibitors.**

| Excipient | Functional Category | $K_i$ (µM)<br>(95% confidence intervals) |
|---|---|---|
| Benzalkonium Chloride | Antimicrobial agent | 62.1 (20.2-187) |
| Butylparaben | Antimicrobial agent | 44.3 (33.8-58.0) |
| D&C Brown No. 1 | Azo dye | 3.11 (2.31-4.20) |
| D&C Orange No. 4 | Azo dye | 2.11 (1.86-2.41) |
| D&C Red No. 33 | Azo dye | 58.1 (49.2-68.7) |
| FD&C Red No. 4 | Azo dye | 8.10 (7.34-8.95) |
| D&C Red No. 6 | Azo dye | 11.3 (7.69-16.7) |
| FD&C Red No. 40 | Azo dye | 2.59 (1.92-3.50) |
| FD&C Yellow No. 6 | Azo dye | 68.4 (58.3-80.4) |
| Naphthol Blue Black | Azo dye | 0.40 (0.32– 0.49) |
| D&C Green No. 5 | Dye | 1.54 (1.18-2.01) |
| D&C Red No. 27 | Dye | 1.0 (0.45-1.31) |
| D&C Red No. 28 | Dye | 1.0 (0.65-1.57) |
| EXT. D&C Yellow No. 7 | Dye | 24.5 (21.9-27.5) |
| FD&C Blue No. 1 | Dye | 13.0 (11.7-14.4) |
| FD&C Red No. 3 | Dye | 0.88 (0.69-1.11) |
| Guinea Green B | Dye | 0.77 (0.64-0.92) |
| Light green CF Yellowish | Dye | 0.80 (0.73-0.89) |
| Ponceau Xylidine | Dye | 3.45 (3.04-3.92) |
| Rhodamine B | Dye | 44.4 (36.4-54.0) |
| Neohesperidin Dihydrochalone | Flavoring agent | 20.1 (16.2-24.8) |
| Docusate Sodium | Surfactant | 2.76 (1.41-5.42) |
| Sodium Lauryl Sulfate | Surfactant | 1.98 (1.37-2.85) |
| Sucrose Monolaurate | Surfactant | 47.7 (36.8-61.9) |

**Table S2. Aggregation analysis for OATP2B1 transport inhibitors.**

| Excipient | $K_i$ (µM) | Aggregation |
|---|---|---|
| Naphthol Blue Black | 0.40 | No Aggregation @ 5 µM |
| Light green CF Yellowish | 0.80 | No Aggregation @ 200 µM |
| FD&C Red No. 3 | 0.88 | No Aggregation @ 500 µM |
| D&C Red No. 28 | 1.0 | No Aggregation @ 10 µM |
| Sodium Lauryl Sulfate | 1.98 | No Aggregation @ 50 µM |
| D&C Orange No. 4 | 2.11 | No Aggregation @ 100 µM |
| FD&C Red No. 40 | 2.59 | No Aggregation @ 500 µM |
| D&C Brown No. 1 | 3.11 | No Aggregation @ 50 µM |
| FD&C Blue No. 1 | 13.0 | No Aggregation @ 200 µM |
| Neohesperidin Dihydrochalone | 20.1 | No Aggregation @ 200 µM |

**Table S3. Physicochemical features of inhibitors versus non-inhibitors.**

| | Inhibitors | | Non-inhibitors | | *p*-value |
|---|---|---|---|---|---|
| | **Mean** | **SD** | **Mean** | **SD** | |
| MW | 504.0 | 181.0 | 216.0 | 157.0 | 4E-08 |
| HA | 31.6 | 9.6 | 14.7 | 10.9 | 5E-09 |
| MV | 401.0 | 124.0 | 195.0 | 132.0 | 1E-08 |
| RB | 7.0 | 4.7 | 4.0 | 4.3 | 1E-03 |
| HBD | 2.9 | 2.0 | 2.9 | 2.9 | ns |
| HBA | 8.2 | 3.5 | 5.2 | 4.3 | 1E-03 |
| SLogP | 6.5 | 2.6 | 1.1 | 2.6 | 7E-11 |
| TPSA | 133.0 | 61.5 | 90.5 | 72.6 | 5E-03 |

Acronyms: MW, molecular weight; HA, number of heavy atoms; RB, number of rotatable bonds; MV, molecular volume; HBD, number of hydrogen bond donors; HBA, number of hydrogen bond acceptors; SLogP, Crippen's log of the octanol/water partition coefficient (including implicit hydrogens); and TPSA, total polar surface area.

**Table S4. Pharmacokinetics of fexofenadine with increasing doses of FD&C Red No. 40.**

| Parameter | Fexofenadine + vehicle | Fexofenadine + 2.5 mg/kg FD&C Red No. 40 | Fexofenadine + 25 mg/kg FD&C Red No. 40 |
|---|---|---|---|
| N | 8 | 4 | 9 |
| $C_{max}$ (nmol/L) | 107 ± 38.9 | 111 ± 13.4 (a) | 70.0 ± 33.26 (b,c) |
| $AUC_{0-360}$ (min*µmol/L) | 29.0 ± 10 | 27.1 ± 3.5 (d) | 15.4 ± 5.98 (e, f) |
| $AUC_{0-480}$ (min*µmol/L)[#] | 40.4 ± 11.4 | 33.2 ± 4.15 (g) | 18.5 ± 5.31 (h, i) |
| $T_{max}$ (min) | 90.0 (30.0-240) | 90.0 (60.0 - 120) | 120 (30.0 - 240) |

Statistics (Tukey's multiple comparisons test): (a) p=0.9822, compared to fexofenadine group; (b) p=0.0815, compared to fexofenadine group; (c) p=0.1304, compared to fexofenadine + 2.5 mg/kg FD&C Red No. 40 group; (d) p=0.8243, compared to fexofenadine group; (e) p=0.0026, compared to fexofenadine group; (f) p=0.0469, compared to fexofenadine + 2.5 mg/kg FD&C Red No. 40 group; (g) p=0.3731, compared to fexofenadine group; (h) p=0.0014, compared to fexofenadine group; (i) p=0.0304, compared to fexofenadine + 2.5 mg/kg FD&C Red No. 40 group. [#]The N for fexofenadine group, fexofenadine + 2.5 mg/kg FD&C Red No. 40, and fexofenadine + 25 mg/kg FD&C Red No. 40 is 5, 4 and 6, respectively.

**Table S5. Inhibitor constant ($K_i$) values for azo dyes and their metabolites.**

| Excipient azo dyes | $K_i$ (µM) | Azo dye metabolite | Max screening concentration (µM) | $K_i$ (µM) |
|---|---|---|---|---|
| D&C Brown No. 1 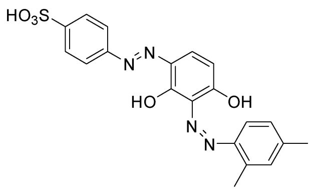 | 3.11 | Sulfanilic acid 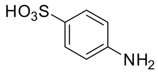 | 200 | >200 |
| | | 2,4-Diaminobenzene-1,3-diol 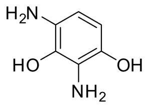 | 200 | >200 |
| | | 2,4-Dimethylaniline 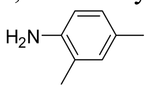 | 200 | >200 |
| D&C Orange No. 4 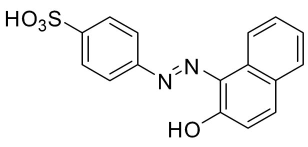 | 2.11 | 1-Amino-2-naphthol 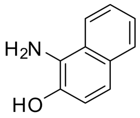 | 200 | 62.5 |
| | | Sulfanilic acid 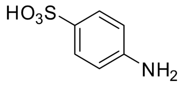 | 200 | >200 |
| D&C Red No. 33 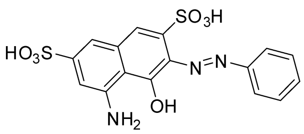 | 58.1 | 3,5-Diamino-4-hydroxy-naphthalene-2,7-disulfonic acid 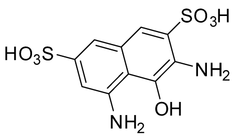 | 50 | >50 |

| | | | | |
|---|---|---|---|---|
| | | Aniline 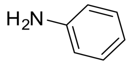 | 200 | >200 |

**Table S5. Inhibitor constant ($K_i$) values for azo dyes and their metabolites continued.**

| Excipient azo dyes | $K_i$ (µM) | Azo dye metabolite | Max screening concentration (µM) | $K_i$ (µM) |
|---|---|---|---|---|
| D&C Red No. 6 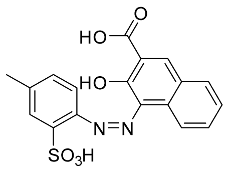 | 11.3 | 4-Amino-3-hydroxy-[2]naphthoic acid 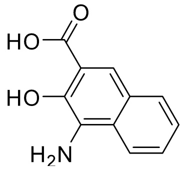 | 200 | >200 |
| | | 4-Aminotoluene-3-sulfonic acid 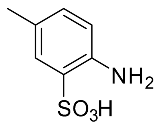 | 200 | >200 |
| FD&C Red No. 4 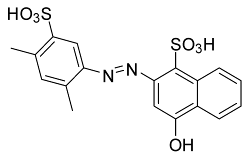 | 8.10 | 5-Amino-2,4-xylenesulfonic acid 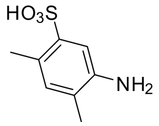 | 125 | >125 |
| | | Unknown metabolite | NA* | NA* |
| FD&C Red No. 40 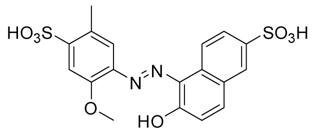 | 2.59 | 4-Amino-5-methoxy-2-methyl-benzenesulfonic acid 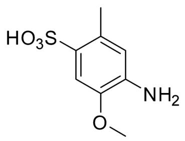 | 50 | >50 |

| | | | | |
|---|---|---|---|---|
| | | 5-Amino-6-hydroxy-2-naphthalene-2-sulfonic acid 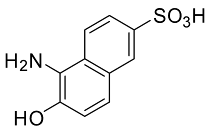 | 200 | >200 |

**Table S5. Inhibitor constant ($K_i$) values for azo dyes and their metabolites continued.**

| Excipient azo dyes | $K_i$ (µM) | Azo dye metabolite | Max screening concentration (µM) | $K_i$ (µM) |
|---|---|---|---|---|
| FD&C Yellow No. 6 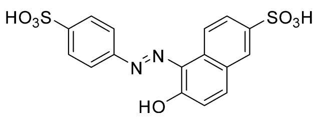 | 68.4 | Sulfanilic acid 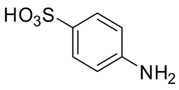 | 200 | >200 |
| | | 5-Amino-6-hydroxy-2-naphthalene-2-sulfonic acid 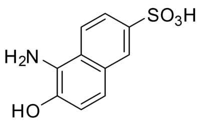 | 200 | >200 |
| Naphthol Blue Black 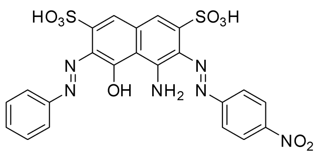 | 0.40 | 4-Nitroaniline 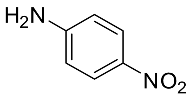 | 200 | >200 |
| | | Aniline 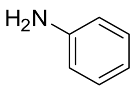 | 200 | >200 |
| | | Unknown metabolite | NA* | NA* |

* NA, not available

**Table S6. Estimated daily intake of FD&C dyes.**

| Dye | Estimated daily intake (mg)* | $K_i$ (μM) | Intestinal $C_{max}$ (μM) | $C_{max}/K_i$ |
|---|---|---|---|---|
| FD&C Red No. 40 | 24.80 | 2.59 | 220 | 84.9 |
| FD&C Yellow No. 6 | 16.70 | 68.4 | 165 | 2.41 |

**Table S7. Estimated daily intake of excipients based on maximum allowable amount.**

| Excipient | Max allowable amount | mg/day[#] | Intestinal $C_{max}$ (µM)[##] | $K_i$ (µM) | $C_{max}/K_i$ |
|---|---|---|---|---|---|
| FD&C Red No. 40 | 7 mg/kg bw* | 490 | 3950 | 2.59 | 1520 |
| FD&C Blue No. 1 | 6 mg/kg bw* | 420 | 2117 | 13.0 | 163 |
| Sodium Lauryl Sulfate | 25 mg/L** | NA | 87 | 1.98 | 44 |
| FD&C Red No. 3 | 0.1 mg/kg bw* | 7 | 32 | 0.88 | 36 |
| FD&C Yellow No. 6 | 4 mg/kg bw* | 280 | 2480 | 68.4 | 36 |
| Docusate Sodium | 0.1 mg/kg bw* | 7 | 63 | 2.76 | 23 |

[*] ADI (Accepted daily intake) (http://apps.who.int/food-additives-contaminants-jecfa-database/search.aspx)
[**] CFR 21 (https://www.accessdata.fda.gov/scripts/cdrh/cfdocs/cfcfr/CFRSearch.cfm?fr=172.822)
[#] Assuming adult body weight of 70 kg
[##] Assuming excipient dissolved in 250 mL liquid
NA, not available

**Supplementary References**


34. J. J. Irwin, J. Pottel, L. Zou, H. Wen, S. Zuk, X. Zhang, T. Sterling, B. K. Shoichet, R. Lionberger, K. M. Giacomini, A Molecular Basis for Innovation in Drug Excipients. *Clin. Pharmacol. Ther.* **101**, 320–323 (2017).

35. A. R. Erdman, L. M. Mangravite, T. J. Urban, L. L. Lagpacan, R. A. Castro, M. de la Cruz, W. Chan, C. C. Huang, S. J. Johns, M. Kawamoto, D. Stryke, T. R. Taylor, E. J. Carlson, T. E. Ferrin, C. M. Brett, E. G. Burchard, K. M. Giacomini, The human organic anion transporter 3 (OAT3; SLC22A8): genetic variation and functional genomics. *Am. J. Physiol. Renal Physiol.* **290**, F905–12 (2006).

36. J. H. Zhang, T. D. Chung, K. R. Oldenburg, A Simple Statistical Parameter for Use in Evaluation and Validation of High Throughput Screening Assays. *J. Biomol. Screen.* **4**, 67–73 (1999).

37. International Transporter Consortium, K. M. Giacomini, S.-M. Huang, D. J. Tweedie, L. Z. Benet, K. L. R. Brouwer, X. Chu, A. Dahlin, R. Evers, V. Fischer, K. M. Hillgren, K. A. Hoffmaster, T. Ishikawa, D. Keppler, R. B. Kim, C. A. Lee, M. Niemi, J. W. Polli, Y. Sugiyama, P. W. Swaan, J. A. Ware, S. H. Wright, S. W. Yee, M. J. Zamek-Gliszczynski, L. Zhang, Membrane transporters in drug development. *Nat. Rev. Drug Discov.* **9**, 215–236 (2010).

38. L. Zhang, Y. D. Zhang, J. M. Strong, K. S. Reynolds, S.-M. Huang, A regulatory viewpoint on transporter-based drug interactions. *Xenobiotica*. **38**, 709–724 (2008).

39. K. E. D. Coan, B. K. Shoichet, Stoichiometry and physical chemistry of promiscuous aggregate-based inhibitors. *J. Am. Chem. Soc.* **130**, 9606–9612 (2008).

40. S. Kim, P. A. Thiessen, E. E. Bolton, J. Chen, G. Fu, A. Gindulyte, L. Han, J. He, S. He, B. A. Shoemaker, J. Wang, B. Yu, J. Zhang, S. H. Bryant, PubChem Substance and Compound databases. *Nucleic Acids Res.* **44**, D1202–13 (2016).

41. M. Sud, MayaChemTools: An Open Source Package for Computational Drug Discovery. *J. Chem. Inf. Model.* **56**, 2292–2297 (2016).

42. L. Kotthoff, *fselector* (Github; https://github.com/larskotthoff/fselector).

43. J. Versalovic, T. Koeuth, J. R. Lupski, Distribution of repetitive DNA sequences in eubacteria and application to fingerprinting of bacterial genomes. *Nucleic Acids Res.* **19**, 6823–6831 (1991).

44. S. Turner, K. M. Pryer, V. P. Miao, J. D. Palmer, Investigating deep phylogenetic relationships among cyanobacteria and plastids by small subunit rRNA sequence analysis. *J. Eukaryot. Microbiol.* **46**, 327–338 (1999).
